# Functional study of *Leishmania braziliensis* protein arginine methyltransferases (PRMTs) reveals that PRMT1 and PRMT5 are required for macrophage infection

**DOI:** 10.1101/2021.09.22.461376

**Authors:** Lucas Lorenzon, José C. Quilles Junior, Gustavo Daniel Campagnaro, Leticia Almeida, Flavio Protasio Veras, Rubens D. M. Magalhães, Juliana Alcoforado Diniz, Tiago R. Ferreira, Angela K. Cruz

**Affiliations:** Department of Cell and Molecular Biology, Ribeirão Preto Medical School, University of São Paulo, Ribeirão Preto, São Paulo, Brazil; Department of Pharmacology, Ribeirão Preto Medical School, University of São Paulo, Ribeirão Preto, São Paulo, Brazil; Laboratory of Parasitic Diseases, National Institutes of Allergy and Infectious Diseases, National Institutes of Health, Bethesda, MD, USA

## Abstract

In trypanosomatids, regulation of gene expression occurs mainly at the posttranscriptional level, and RNA-binding proteins (RBPs) are key players in determining the fates of transcripts. RBPs are major targets of protein arginine methyltransferases (PRMTs), which posttranslationally regulate the RNA-binding capacity and other macromolecular interactions of RBPs by transferring methyl groups to protein arginine residues. Herein, we present the results of a study that functionally characterized the five predicted PRMTs in *Leishmania braziliensis* by gene knockout and endogenous protein HA tagging using CRISPR/Cas9 gene editing. We report that arginine methylation profiles vary among *Leishmania* species and that target protein methylation changes across different *L. braziliensis* life cycle stages, with higher PRMT expression in the promastigote stages than in the axenic amastigote stage. Knockout of some of the *L. braziliensis* PRMTs led to significant changes in global arginine methylation patterns without affecting promastigote axenic growth. Deletion of either PRMT1 or PRMT3 disrupted most type I PRMT activity, resulting in a global increase in monomethyl arginine (MMA) levels, which is mainly catalyzed by PRMT7. Putative targets and/or PRMT-interacting proteins were identified by coimmunoprecipitation using HA-tagged PRMTs, revealing a network of target RBPs and suggesting functional interactions between them and a relevant participation in epigenetic control of gene expression. Finally, we demonstrate that *L. braziliensis* PRMT1 and PRMT5 are required for efficient macrophage infection *in vitro*, and that in the absence of PRMT1 and PRMT5, axenic amastigote proliferation is impaired. The results indicate that arginine methylation is modulated across life cycle stages in *L. braziliensis* and show possible functional overlap and cooperation among the different PRMTs in targeting proteins. Overall, our data suggest important regulatory roles of these proteins throughout the *L. braziliensis* life cycle, showing that arginine methylation is important for parasite-host cell interactions.

## 1. Introduction

Posttranslational modifications (PTMs) are known to regulate protein function in multiple processes, including the coordination of regulatory gene expression networks. Protein arginine methylation is a common PTM known to affect RNA-binding protein (RBP) activity (1-3) in several eukaryotes, including early-branching protozoa. The enzymes responsible for catalyzing the transfer of a methyl group from S-adenosyl-methionine to the terminal nitrogen of a peptidyl arginine residue are called **<u>P</u>**rotein a**<u>R</u>**ginine **<u>M</u>**ethyl**<u>T</u>**ransferases (PRMTs); these enzymes compose a nine-member family of proteins and have been identified in a large number of eukaryotes (4). PRMTs are involved in many cellular functions, from transcription regulation to RNA splicing, DNA repair, cell cycle and signal transduction (1). There is clear evidence that nucleic acid-binding proteins are the major substrates of PRMTs and that RBPs account for a large proportion of these PRMT substrates (5, 6). PRMTs are divided into three major types, all of which can catalyze ω-N^G^-monomethylarginine (MMA) as an initial methylation step; type I PRMTs catalyze the formation of asymmetric ω-N^G^-dimethylarginine (ADMA) as a final product, type II PRMTs produce symmetric ω-N^G^-dimethylarginine (SDMA) and type III PRMTs catalyze MMA alone. It is widely accepted that PRMT activity does not alter the charge of the target arginine residue, but an increase in hydrophobicity and mass may affect the hydrogen bonding capability of the target residue, thus positively or negatively modifying macromolecular interactions (7, 8).

Recent descriptions of PRMT (dys)function in many cellular mechanisms, including cancer progression and inflammatory responses, have emphasized the need for a better understanding of the role of arginine methylation in health and disease (9, 10). Notably, the importance of PRMTs in infectious diseases remains largely unknown; in particular, intracellular eukaryotic pathogens also express PRMT homologous genes, which may serve as putative drug targets for antiparasitic therapies.

Trypanosomatids, such as *Trypanosoma* spp. and *Leishmania* spp., are early-branching, medically relevant protozoa that carry five different classic PRMT genes (PRMT1, PRMT3, PRMT5, PRMT6 and PRMT7). In fact, these are the only protozoan parasites known thus far to harbor type I, II and III PRMTs, which suggests that arginine methylation may play a critical role during the life cycles of these protozoa (11, 12).

In addition to their medical relevance, trypanosomatids are unique eukaryotes because of their particular genetic organization that regulates gene expression, and this regulation of gene expression is mostly dependent on posttranscriptional regulatory mechanisms (13-16). Within the last two decades, thorough investigations of PRMTs and arginine methylation have been conducted in *Trypanosoma brucei*. Four active PRMT enzymes, namely, *Tb*PRMT1^ENZ^, *Tb*PRMT5, *Tb*PRMT6, and *Tb*PRMT7, and one prozyme (*Tb*PRMT1^PRO^), previously known as *Tb*PRMT3, have been characterized in *T. brucei*. Furthermore, fundamental knowledge about the enzymatic activities, effects of knockdown or knockout, functional interplay and substrate specificities of these enzymes has been obtained (11, 17-25).

Comparatively, much less is known about *Leishmania* PRMTs. *Leishmania* parasites are responsible for approximately 12 million cases of leishmaniasis worldwide; leishmaniasis is a human disease distributed in more than 80 countries in the tropical and subtropical regions of the globe that causes between 20 and 40 thousand deaths per year (26). To date, of the five classic PRMT homologs present in the *Leishmania* genome, only studies on *Leishmania major* PRMT7 have been reported (12). We have previously shown that the *Lmj*PRMT7 levels are inversely correlated with the pathogenicity of *L. major in vitro* and *in vivo* (27, 28). In addition, Ferreira and coworkers showed that *Leishmania* PRMT7 acts as an epigenetic regulator of mRNA metabolism and provided mechanistic insight into the functional regulation of RBPs by methylation (29). Two of the identified *Lmj*PRMT7 substrates, Alba3 and RBP16, were shown to be affected by this posttranslational modification in different ways. A global monomethyl arginine proteomic analysis revealed that approximately 3% of *L. major* proteins, including several RNA-binding proteins, have at least one methylated arginine residue, shedding light on the importance of this posttranslational modification for the parasite (29).

Given the relevance of improving the understanding of eukaryotic pathogens and PRMT function and the lack of information about *Leishmania* PRMTs, we present the first functional investigation of all five canonical PRMTs in *Leishmania braziliensis*, the main causative agent of tegumentary leishmaniasis in South America. We observed that although the three types of arginine methylation are observed in the three main biological forms of *L. braziliensis*, MMA is most likely the most common type. Immunoprecipitation assays revealed that R-methylated proteins are distributed in all cellular compartments, that RBPs are well represented among *Lb*PRMT targets and that some of the *Lb*PRMTs may associate with each other in protein complexes. Moreover, our study reveals that the expression of both type I *Lb*PRMT1 and type II *Lb*PRMT5 is critical for establishing macrophage infection *in vitro*, suggesting that both SDMA and ADMA arginine methylation are important for the development of *bona fide* infectious *L. braziliensis*.

## 2. Results

### Conservation and divergence among PRMTs

Global protein sequence alignments for each of the PRMTs were conducted, and these analyses were particularly focused on key motifs in the methyltransferase domain (6). We compared the protein sequences from *L. braziliensis* with their corresponding orthologs in *T. brucei* and humans. *Lb*PRMT1 had 70.4% and 49% identity with *Tb*PRMT1^ENZ^ and *Hs*PRMT1, respectively; *Lb*PRMT3 had 46.4% and 24.2% identity with these orthologs, respectively; *Lb*PRMT5 had 29.4% and 17.7% identity with these orthologs, respectively; *Lb*PRMT6 had 46.6% and 24.9% identity with these orthologs, respectively; and *Lb*PRMT7 had 50.7% and 21.5% identity with these orthologs, respectively. Overall, most of the six signature domains were conserved between the three species, but Motif III was particularly divergent when trypanosomatid PRMTs were compared with their human orthologs. Interestingly, both *Tb*PRMT1^PRO^ and *Lb*PRMT3 harbored mutations in the THW motif, but the Double E loop was altered only in *Tb*PRMT1^PRO^ (Fig. **1**).

**Figure 1.**
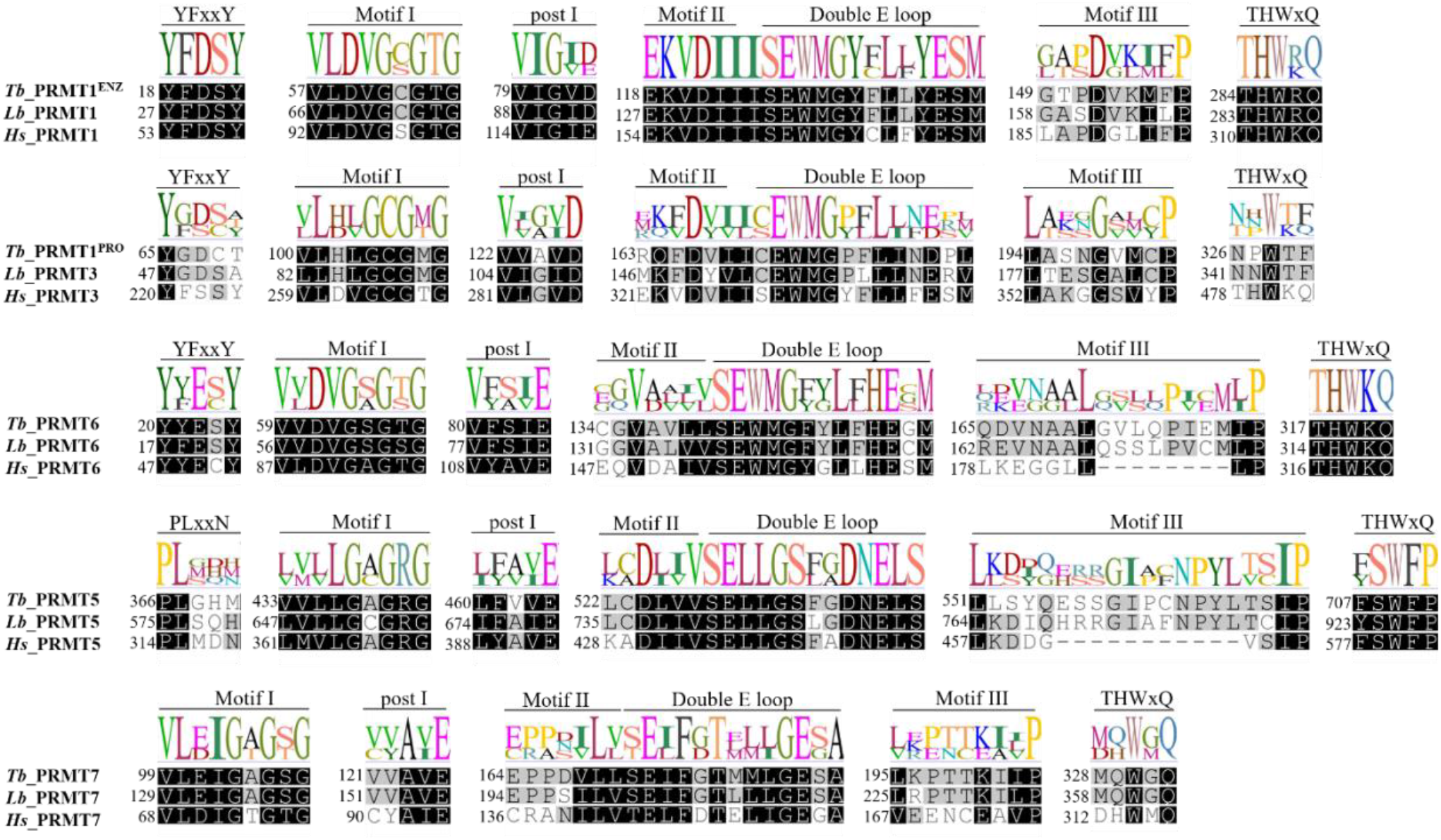
Global sequence alignment of PRMTs from *L. braziliensis, T. brucei* and *Homo sapiens*. The protein sequences were aligned using Geneious v6.18 software with Clustal Omega. The key motifs of the conserved PRMT methyltransferase domain are indicated. Residue shading: gray background: 60–80% similar; black background: 80– 100% similar. The amino acid sequence logo is shown on the top of each alignment representing motif sequence conservation.

We also compared the sequences of PRMTs from parasitic and nonparasitic kinetoplastida. We observed that the motifs were highly conserved, with few exceptions. Interestingly, even the free-living kinetoplastid *Bodo saltans*, the closest relative of representative trypanosomatid parasites, contained the same conserved signatures (Fig. **S1**), highlighting the conservation of these regions during evolution.

### Differential arginine methylation patterns in distinct *Leishmania* species and parasite life cycle stages

To evaluate arginine methylation in *Leishmania*, we used polyclonal antibodies that recognize each of the three major types of R-methylation (MMA, SDMA and ADMA) (Fig. **2a**). As observed by western blotting, the three types of R-methylation were present in the promastigotes of different species of *Leishmania* (Fig. **2b, 2c** and **2d**), with evidence of slightly distinct R-methylation profiles, especially the MMA profiles, among the analyzed species. However, comparative quantitative analysis among the three different R-methylation types is not possible with this approach due to the differences in antibody origin and staining patterns used for each type. Next, we evaluated the profiles of R-methylated proteins in *L. braziliensis* throughout the progression of its life cycle, namely, in procyclic and metacyclic promastigotes and in axenic amastigotes (Fig. **2e, 2f** and **2g**). R-methylation was present in all the observed life cycle stages, with some detectable differences in the profiles, especially in the MMA and ADMA profiles, among these stages; most strikingly, a band of ∼33 kDa, containing ADMA, was enriched in the metacyclic promastigote stage compared to the other two stages. Moreover, as a general rule, amastigotes seemed to harbor a lower level of protein R-methylation than promastigotes (Fig. **2f** and **2h**).

**Figure 2.**
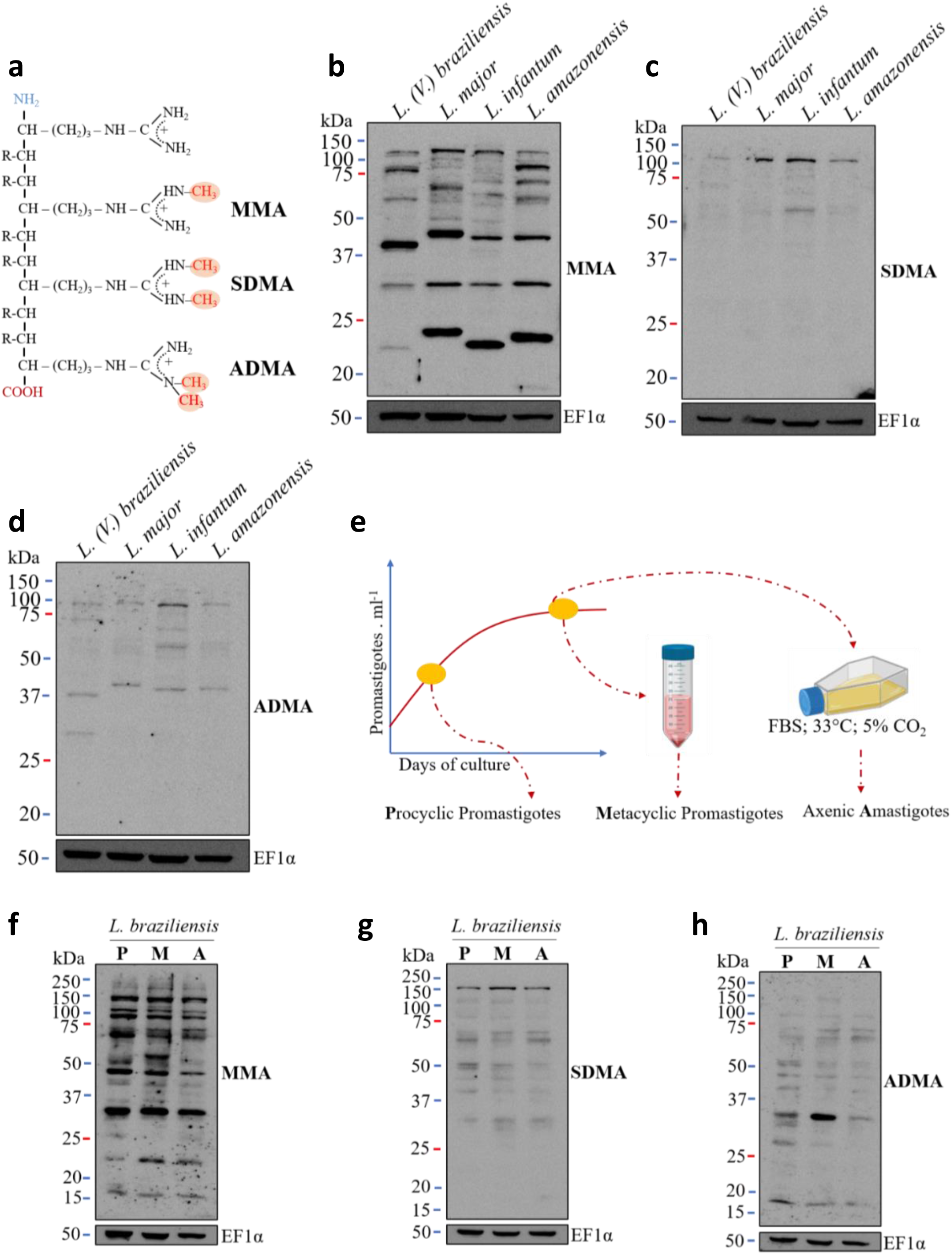
Arginine methylation is present in different *Leishmania* species and stages of the *Leishmania* life cycle. **a**. Schematic representation of the three major types of protein arginine methylation catalyzed by PRMTs. **b, c** and **d**. The profiles of monomethylated (MMA), symmetrically dimethylated (SDMA) and asymmetrically dimethylated (ADMA) arginine in different species of *Leishmania* were determined by western blotting. **e**. A schematic representation of the protocol used to obtain samples from three distinct stages of the *L. braziliensis* life cycle. **f, g** and **h**. The profiles of MMA, SDMA and ADMA of distinct stages of the *L. braziliensis* life cycle; (P) promastigotes, (M) metacyclic promastigotes and (A) amastigotes. The loading control for all the western blots (Panels **b, c, d, f, g** and **h**) is shown on the lower blots using the anti-EF1alpha antibody.

### PRMTs are mainly cytoplasmic proteins that are preferentially expressed in the promastigote stages

The differences in the R-methylation profiles observed in *L. braziliensis* promastigotes and axenic amastigotes suggested that this PTM might be important for parasite development. Thus, we evaluated the expression profiles of the five putative *Lb*PRMTs in the three main stages of the life cycle. Due to the difficulties associated with genetically manipulating *Leishmania* spp. via classical homologous recombination techniques, we repurposed the CRISPR/Cas9 system previously developed by Beneke et al. (28) and generated an *L. braziliensis* strain expressing both Cas9 and T7 RNA polymerase, named pT007. The expression of Cas9 was confirmed by western blotting with antibodies against the Flag tag (Fig. **3a**). Next, each of the PRMT coding sequences was endogenously tagged with three tandem copies of the HA tag at the 5’ end (Fig. **3b**). The correct insertion of the HA tag into both alleles of each targeted locus was confirmed by PCR (Fig. **3c**); the transfectants were named tag1, tag3, tag5, tag6 and tag7 (where the number indicates each of the tagged PRMTs). The expression of the tagged PRMTs (HA-PRMT1, HA-PRMT3, HA-PRMT5, HA-PRMT6 and HA-PRMT7) in the transfectants was confirmed by western blotting using protein extracts from procyclic promastigotes. The expected molecular weight of each PRMT was confirmed (Fig. **3d**). The same *L. braziliensis* transfectants were used to evaluate the expression of the PRMTs across developmental stages (Fig. **3e**). Western blotting results of protein extracts from procyclic and metacyclic promastigotes and axenic amastigotes indicated that the tagged PRMTs were preferentially expressed in promastigotes rather than in amastigotes (Fig. **3e**). Decreased expression in amastigotes was observed for all the PRMTs, but this decrease in expression was particularly evident for PRMT7, which was barely detected in this biological form (Fig. **3e**). We further investigated the subcellular distribution of the PRMTs using the same transfectants. Confocal microscopy analysis revealed differences in the intracellular distribution of the PRMTs: PRMT1, PRMT3 and PRMT5 were detected in both the cytoplasm and nucleus, PRMT6 was observed mainly in the intranuclear space, and PRMT7 was found only in the cytoplasm (Fig. **3f**).

**Figure 3.**
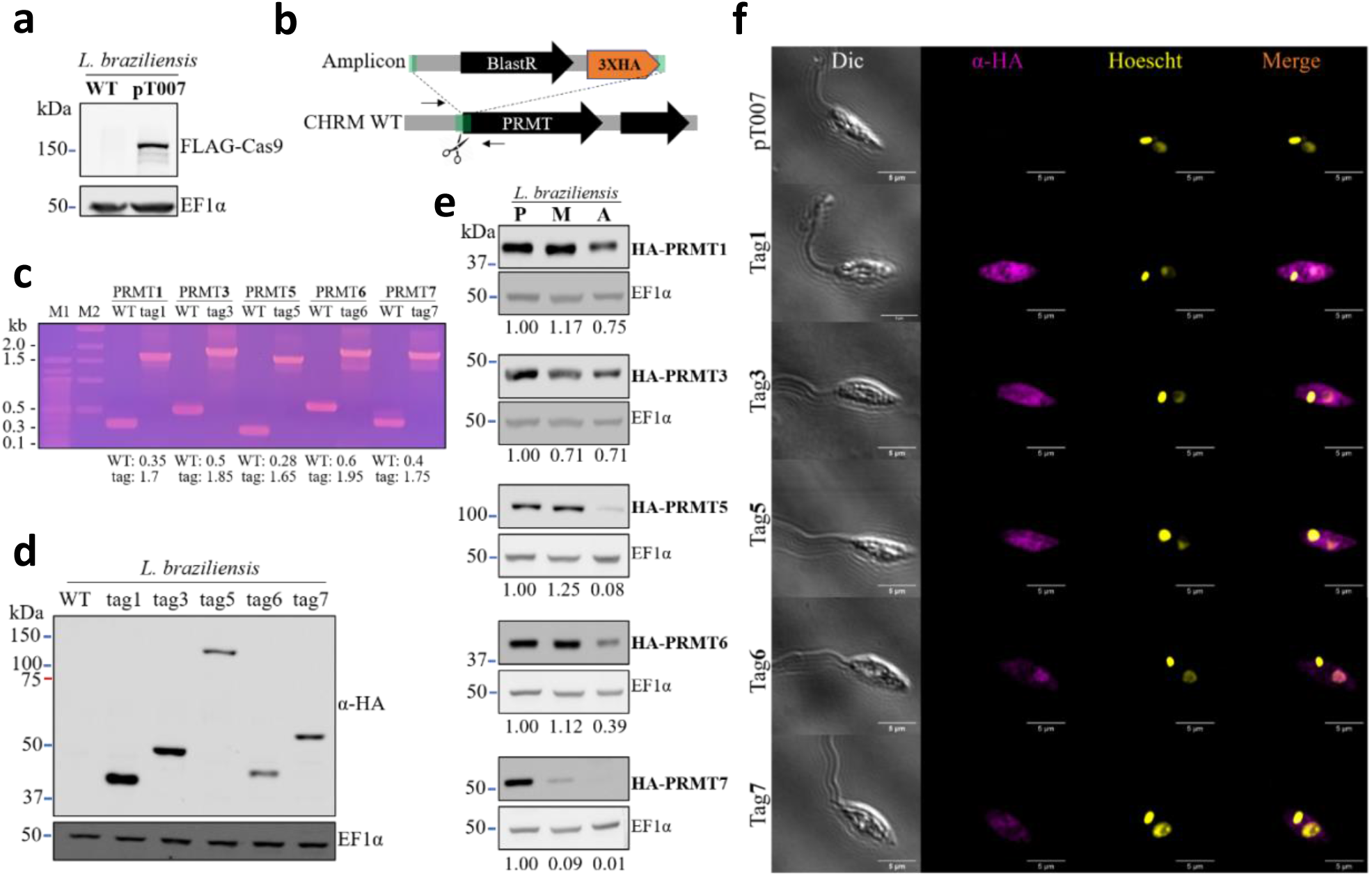
HA-tagged PRMTs are preferentially expressed in promastigotes, and most of them are cytoplasmic. **a**. Expression of FLAG-Cas9 in *L. braziliensis* pT007, as shown by western blotting with anti-FLAG antibodies (Sigma F1804). **b**. Schematic representation of the strategy used for HA tagging of the N-terminus of each PRMT coding gene. BlastR represents the blasticidin resistance gene; the black arrow refers to the PRMT coding sequence in the respective genomic locus (CHRM: chromosome), and the green boxes represent a 30-bp homology to the locus to be edited; the thin black arrows depict the annealing site of the primers used in the diagnostic PCR shown in *c*. **c**. Confirmation of PRMT endogenous HA tagging by PCR analysis and agarose gel (1%) electrophoresis showing the expected amplicon length for each of the tagged PRMTs for wild-type (WT) and tagged PRMTs (tag) in kilobase pairs. M1 and M2 are standard MW markers (100 bp and 1 kb, respectively – both from New England Biolabs). **d**. Expression of HA-tagged PRMTs (tag1 indicates HA-PRMT1; tag3 indicates HA-PRMT3, and so on) in *L. braziliensis* procyclic promastigotes. The blots confirm the expected molecular weight for each PRMT. In the upper panel, an anti-HA antibody (H3663 – Sigma Aldrich) was used. **e**. Evaluation of HA-PRMT expression throughout development; procyclic promastigotes (P), metacyclic promastigotes (M) and axenic amastigotes (A) of *L. braziliensis*. The values below the panels represent the densitometric analysis (PRMT/EF1α). **f**. Confocal microscopy showing the subcellular localization of HA-PRMTs. An anti-HA antibody (H3663 Sigma) was used to detect the HA-PRMTs, while Hoechst 33342 (Invitrogen) was used to detect the mitochondrial and nuclear DNA. EF1alpha was evaluated as a loading control in all the western blotting data shown.

### Knockout of the PRMT1 and PRMT3 genes leads to changes in the arginine methylation profiles

To determine the biological relevance of the *L. braziliensis* PRMTs, we generated null mutants for each of their respective genes. Using established CRISPR/Cas9-expressing pT007 cells, we replaced each PRMT coding sequence with a blasticidin S resistance marker (BlastR, blasticidin deaminase; (30)) (Fig. 4a). Null mutants lacking each PRMT were generated by gene knockout (KO; Δ1, Δ3, Δ5, Δ6 and Δ7, where *Δn* represents the deletion of the *PRMTn* gene), and the results demonstrated the nonessentiality of each individual PRMT for promastigote growth (Supplementary Fig. S2). However, the R-methylation profiles in some of these null mutants were significantly altered (Fig. **4c** – **e**). Interestingly, both the Δ1 and Δ3 transfectants showed markedly increased levels of MMA and a less pronounced, but relevant, increase in the SDMA levels (Fig. **4c** and **4d**, respectively). In contrast, in the absence of either PRMT1 or PRMT3, the levels of ADMA were drastically reduced (Fig. **4e**). To validate these data, we generated constructs to add back each of the PRMTs to the corresponding KO parasites. HA-PRMTs, along with the neomycin phosphotransferase gene as a selectable marker, were inserted into the **S**mall **Su**bunit Ribosomal (SSU rDNA) locus (Fig. **4f**). The integration of linearized DNA constructs into the target locus was confirmed (Supplementary Fig. S2), and the addback transfectants were named Add1, Add3, Add5, Add6 and Add7. The complementation of PRMT1 and PRMT3 clearly restored the MMA, SDMA and ADMA levels to those observed in the parental line (Fig. **4g, h** and **i**).

**Figure 4.**
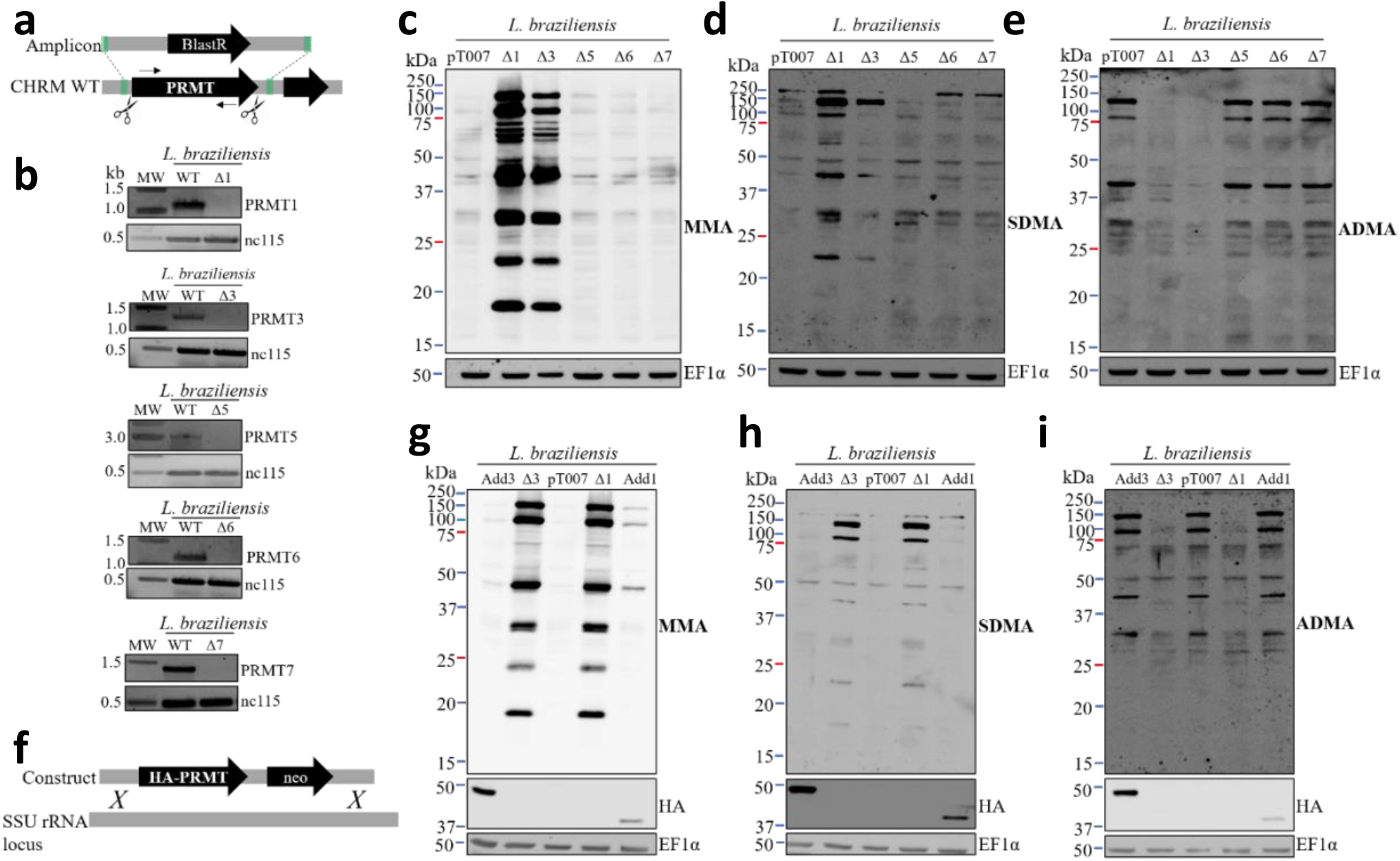
Gene knockout of PRMT1 and PRMT3 leads to changes in arginine methylation profiles. **a**. Schematic representation of the strategy used to generate PRMT-null mutants (black arrows represent the primers used in PCR shown in *b*). **b**. Confirmation of gene deletion in the transfectant clones by PCR using primers that anneal within the PRMT CDS. **c, d** and **e**. Arginine methylation profiles of monomethylated (MMA), symmetrically dimethylated (SDMA) and asymmetrically dimethylated arginine (ADMA) in null mutants (Δ*n* represents PRMT*n* gene deletion) were determined by Western blotting. **f**. Schematic representation of the strategy used to generate PRMT addback parasites. **g, h** and **i**. Arginine methylation profiles of MMA, SDMA and ADMA of addback parasites (Add*n* represents PRMT*n* addback in a Δ*n* background) were determined by Western blotting. CHRM: chromosome; MW: DNA molecular weight marker; nc115 and EF1a were used as loading controls in DNA agarose gels and protein blots, respectively.

### PRMT interactome analysis reveals multiple RNA-binding protein targets and suggests interplay between PRMTs

The differential R-methylation profiles of the mutants lacking PRMT1 and PRMT3 led us to investigate the possible binding partners of each PRMT by coimmunoprecipitation (co-IP) assays. We used HA-tagged proteins to pull down the different PRMTs and the proteins with which they physically interact. The accurate immunoprecipitation of each PRMT was confirmed by Western blotting (Fig. **5a**). A protein band of approximately 47 kDa was detected in all the samples, and this molecular weight was consistent with the coeluted anti-HA antibody heavy chain and with the expected molecular weight of HA-PRMT3. The eluted protein complex was then analyzed by mass spectrometry to identify the proteins. After the removal of contaminating proteins (i.e., keratin, trypsin, serum albumin) and proteins identified in the WT sample (i.e., proteins not related to PRMT interactions), a total of 135 proteins were shown to interact with PRMTs. In summary, 77 proteins were pulled down with PRMT1, 38 with PRMT3, 66 with PRMT5, 44 with PRMT6 and 66 with PRMT7 (Table **S1**). Data analysis showed that most of the possible directly or indirectly interacting proteins were shared among PRMTs, while fewer proteins specifically interacted with only one PRMT (Fig. **5b**). Among the 135 proteins identified, we searched for those with RG motifs, which are known to be preferential sites of R-methylation, and for proteins involved in RNA binding. We identified 20 proteins with one of these characteristics, such as several ATP-dependent RNA helicases and proteins related to both RNA and DNA metabolism (NOL1/NOP2 rRNA methyltransferase, RNA cytidine acetyltransferase, and ruvb-like 1 DNA helicase, among others; Fig. **5c**).

**Figure 5.**
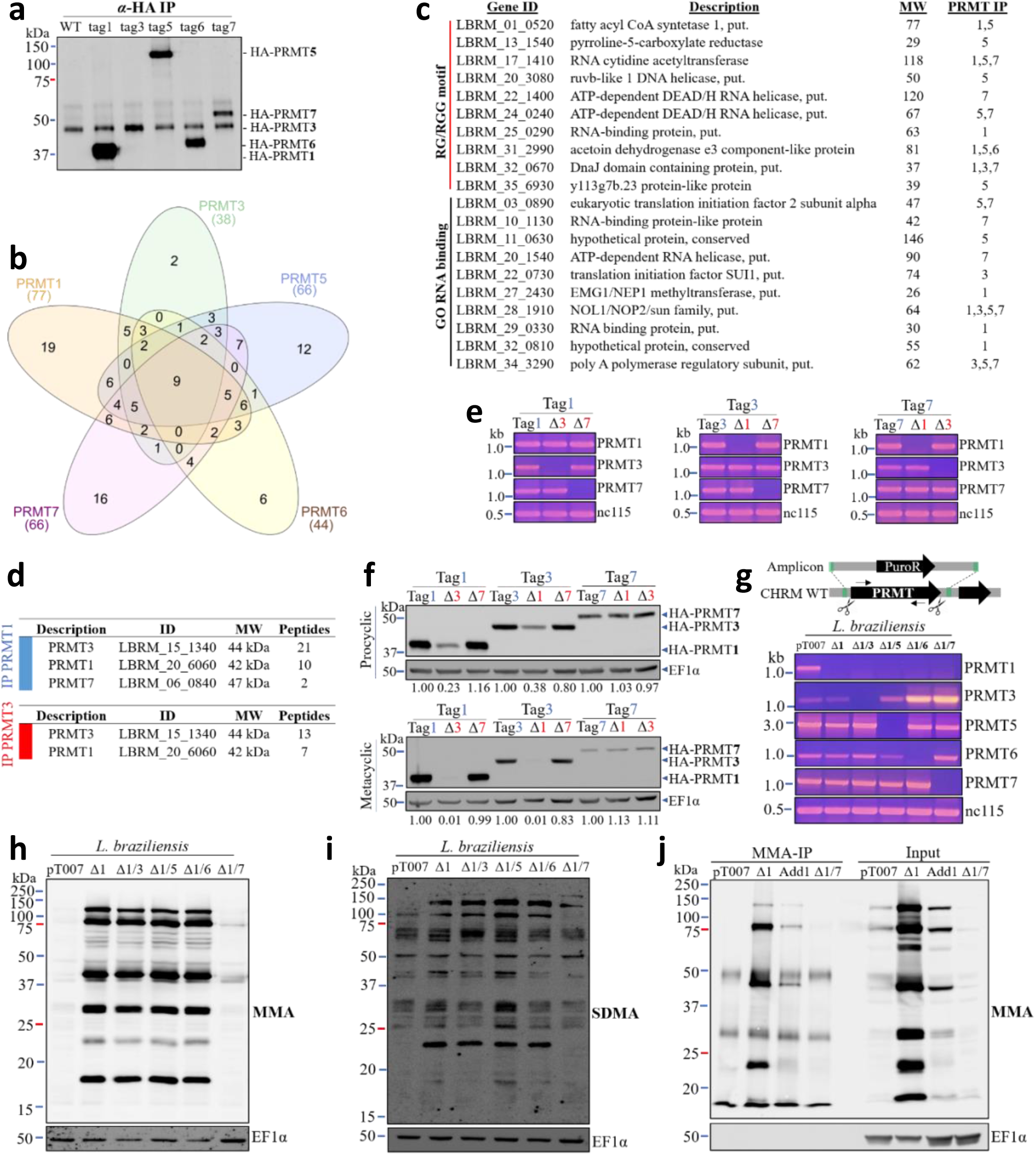
The PRMT interactome has diverse RNA-binding proteins and suggests interplay between PRMTs. **a**. Validation of HA-PRMT immunoprecipitation (IP) by Western blotting with anti-HA antibodies (Tag*n* represents HA-tagged PRMT*n*). **b**. Venn diagram of the proteins co-IP with each of the HA-PRMTs. **c**. List of the PRMT interacting proteins that either have RGG motifs or are associated with GO terms related to RNA binding (MW: molecular weight in kDa, T: true, F: false for the presence of an RG motif or association with an RBP-related GO term) **d**. List of PRMTs interacting with each other as detected in the co-IP assays. **e**. Confirmation of PRMT1/3/7 gene deletion in HA-tagged transfectants by PCR. **f**. Effects of PRMT gene deletion on the expression levels of HA-PRMT1/3/7 as evaluated by Western blotting. **g**. Confirmation of double PRMT gene knockout by PCR; diagram represents the strategy used to generate the double null mutants. **h** and **i**. Arginine methylation profiles of monomethylated (MMA) and symmetrically demethylated (SDMA) in double-null parasites as determined by Western blotting. **j**. Monomethyl arginine (MMA) IP. Western blotting analysis of MMA profiles of proteins immunoprecipitated using an anti-MMA antibody. CHRM WT: wild-type chromosome; nc115 and EF1a were used as loading controls in DNA agarose gels and protein blots, respectively.

Interestingly, two other PRMTs were identified among PRMT1-interacting proteins: PRMT3, the binding protein with most unique peptides identified in this co-IP, and PRMT7 (Fig. **5d**). Similarly, PRMT1 was the second most abundant PRMT3 binding partner (Fig. **5d**). These findings suggest that *L. braziliensis* PRMTs could act as heterocomplexes or even functionally overlap, as already observed in other organisms, including *T. brucei* (25, 31). To investigate this hypothesis, we designed a strategy to knockout the three PRMT genes in the HA-tagged transfectants, aiming to determine whether the absence of either PRMT3 or PRMT7 could lead to changes in the PRMT1 expression levels (similar experiments were performed with the other possible combinations). These tag-knockout combinations were successfully generated (Fig. **5e**). The results clearly showed that knockout of PRMT1 strongly reduced the expression of PRMT3, and the inverse analysis showed similar results, in both procyclic and metacyclic promastigotes (Fig. **5f**). However, neither the knockout of PRMT1 nor PRMT3 altered the expression of PRMT7 (Fig. **5f**).

These results, along with the nonessentiality of the PRMTs (Fig. **4b**), prompted us to generate double knockout mutants and to evaluate the viability of the transfectants. Given the differential R-methylation profiles (Fig. **4c**, 4**d** and **4e**), we knocked out each of the other PRMTs on the PRMT1-null background, generating double null mutants (e.g., Δ1/3 to indicate the knockout of both the PRMT1 and PRMT3 genes). Surprisingly, all the double null transfectants we investigated were viable (Fig. **5g**). We next asked whether the double knockout would also affect the differential R-methylation phenotype. Interestingly, the MMA and SDMA hypermethylation observed in Δ1 (Fig. **4c** and **4d**) was almost completely lost in the absence of PRMT7 (Fig. **5h** and **5i**).

### Monomethyl arginine IP reveals that diverse proteins are hypermethylated in PRMT1-null mutants

Given the arginine methylation profiles observed in the Δ1 and Δ3 parasites (Fig. **4c** and **4d**) together with the finding that PRMT7 is required for MMA accumulation, we aimed to identify proteins that were highly monomethylated in the Δ1 parasite. Using anti-MMA antibodies, we performed an IP experiment to identify MMA proteins in the pT007, Δ1, Add1 and Δ1/Δ7 transfectants (Fig. **5j**). Quantitative analysis of the mass spectrometry data indicated that 32 proteins were at least 2 times more abundant in Δ1 than in all the other transfectants. Among these proteins, RBPs, Rab proteins, one centromere-binding protein (cbf5), one RNA helicase and some uncharacterized proteins were noteworthy (Table **1**), suggesting a role for *L. braziliensis* PRMTs in the control of nucleic acid-binding protein activities.

**Table 1.**
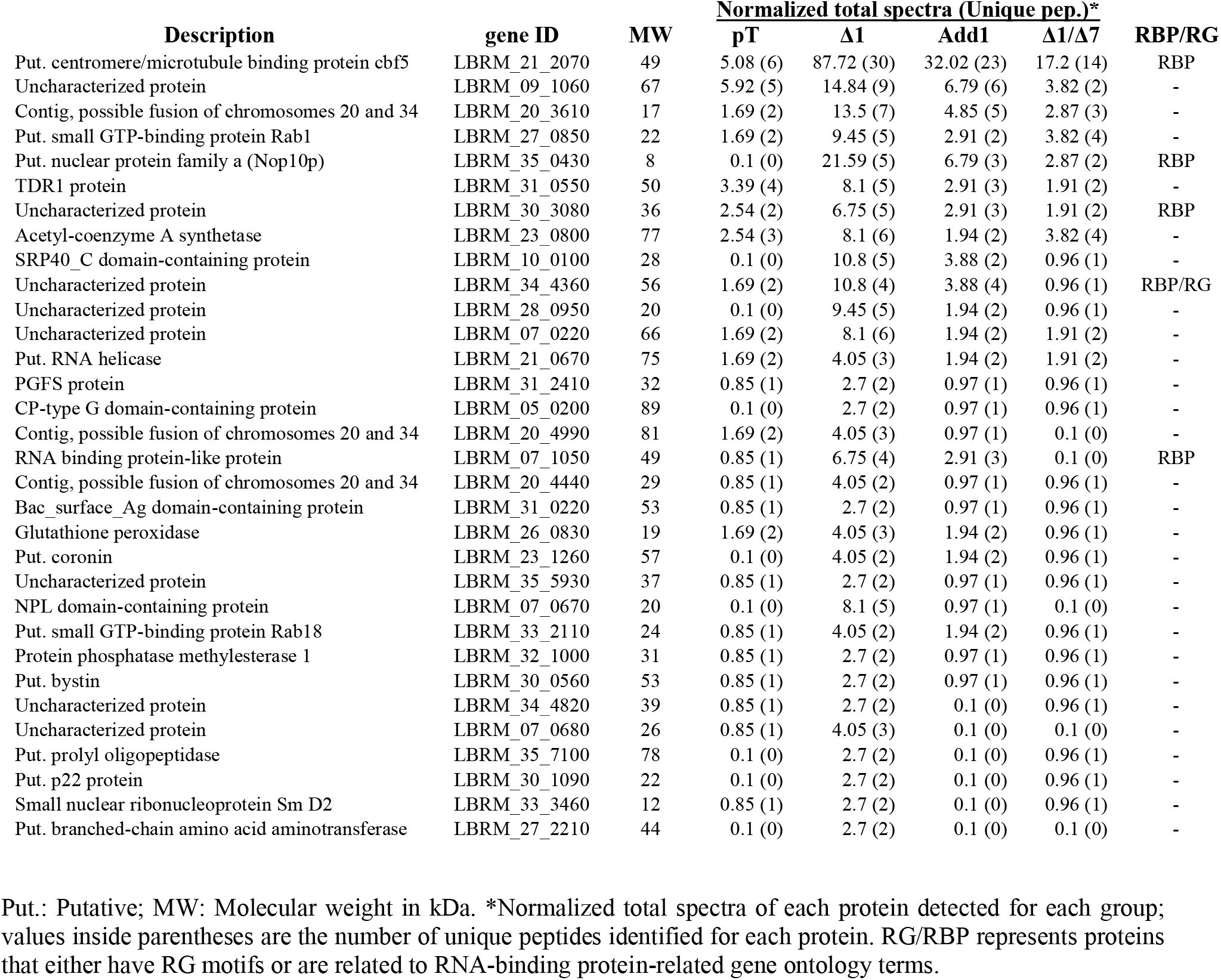
List of the proteins that were at least two times more abundant in Δ1 IP.

Using a complementary MS analysis approach on the same samples, we were also able to identify peptides bearing mono- or dimethylated arginine residues. Within the four samples (pT007, Δ1, Add1 and Δ1/Δ7), we identified MMA peptides corresponding to 38 different proteins; of these proteins, 7 proteins harbored both mono- and dimethylated peptides (Table 2). Additionally, 14 other proteins with DMA peptides were identified. Among these proteins, several putative RNA helicases, RBPs (including RBP16), rRNA methyltransferases and several uncharacterized proteins harboring more than one arginine methylated residue were noteworthy (Table **2**).

**Table 2.**
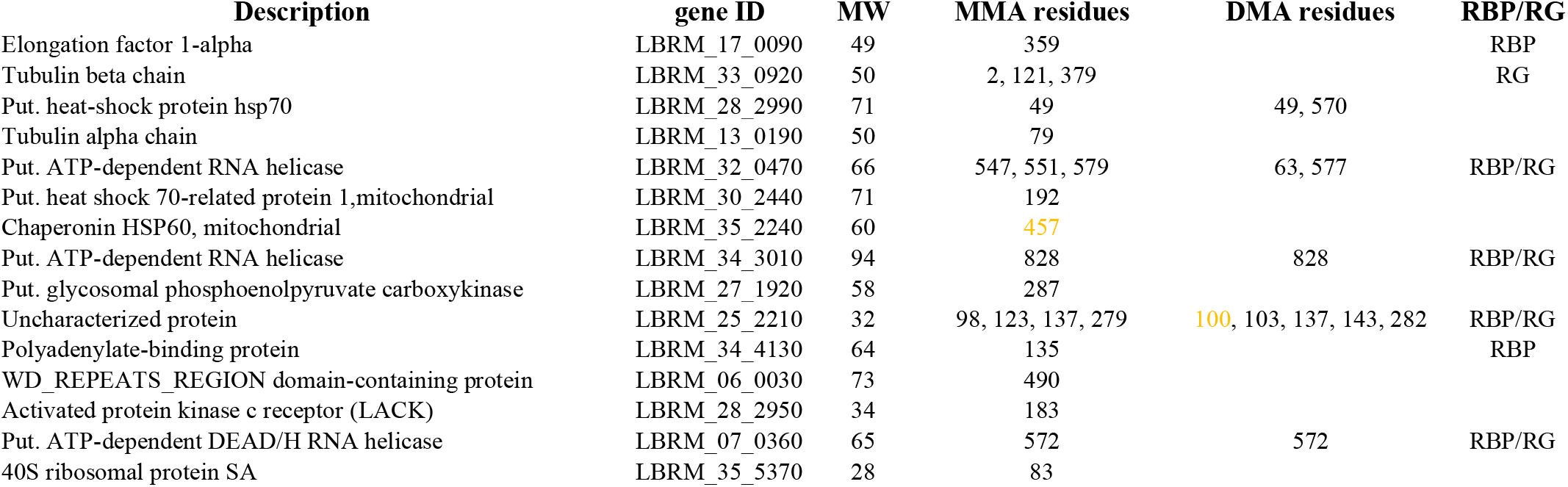
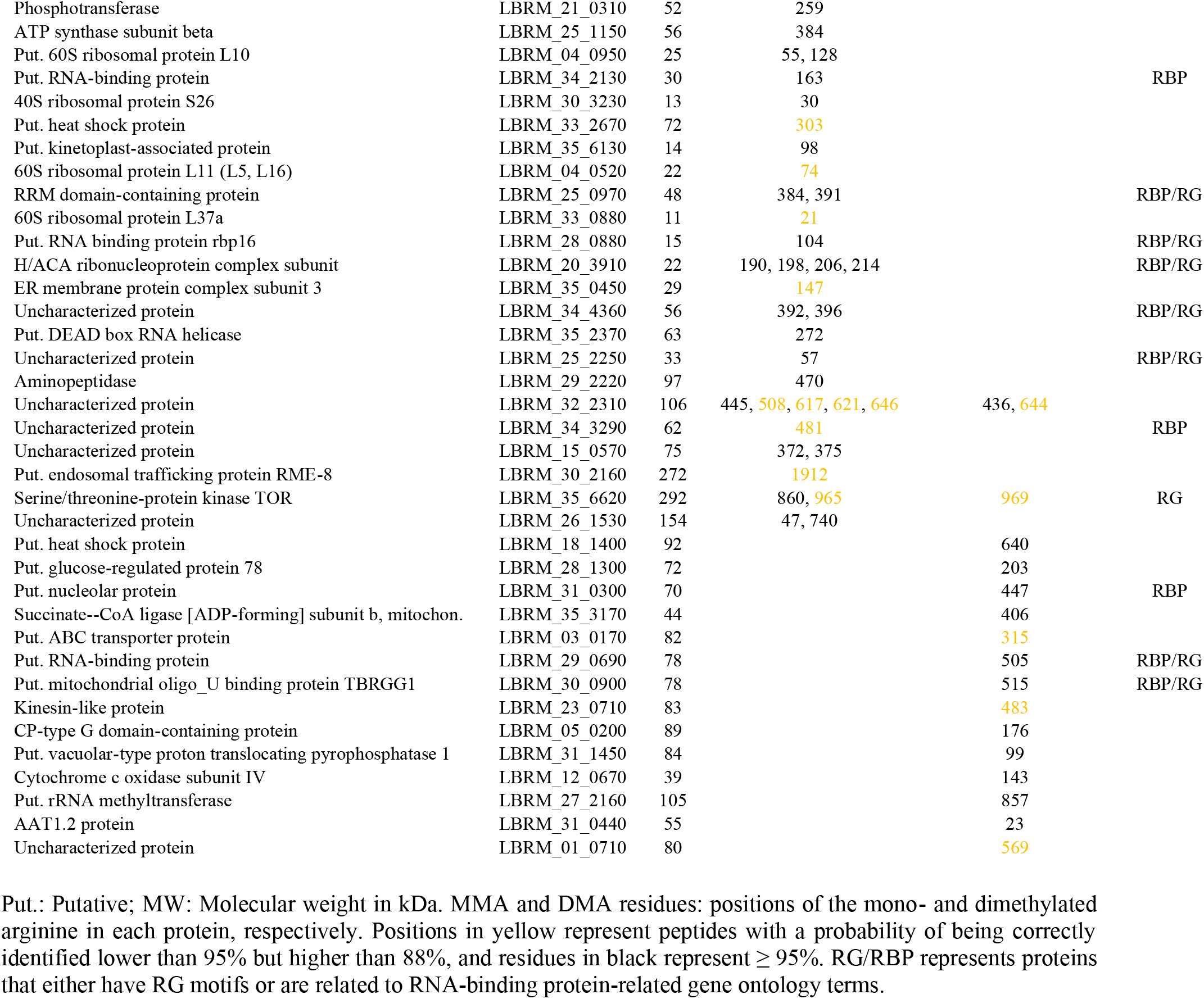
Proteins with R-methyl peptides identified in MMA IP

### PRMT1- and PRMT5-null parasites fail to efficiently infect human and mouse macrophages *in vitro*

To investigate the possible role of *L. braziliensis* PRMTs in parasite-host cell interactions, we first generated *L. braziliensis* parasites expressing both Cas9 and tdTomato (pT_Tomato, Fig. **6a** and **6b**); these parasites allowed the detection and quantitative analysis of *in vitro* infections in an automated manner (Fig. **6c**). Next, we deleted each PRMT gene in the pT_Tomato parasites (Fig. **6d** and **6e**) and used fluorescently labeled null mutants for *in vitro* infection assays. Analyses of the infection profiles and intracellular survival rates were performed in three independent infection assays with the THP-1 monocyte-like cell line. Interestingly, the PRMT1- and PRMT5-null mutants infected both murine and human macrophages at lower ratios than the control parasites (in human macrophages *p* <0.0005 for both PRMT1 and PRMT5, in murine macrophages *p*=0.08 for PRMT1 and *p*=0.0003 for PRMT5). Both the percentage of infected THP-1 phagocytes and the number of intracellular amastigotes were significantly reduced 48 h postinfection with the two mutants (Fig. **6f** and **6g**). Strikingly, there were ∼50-fold and ∼3-fold decreases in the percentage of infected macrophages after infection with the Δ1 and Δ5 mutants, respectively, compared to the percentage of infected macrophages after infection with the control parasites (Fig. **6f**). To validate these data, we generated addback parasites by returning the PRMT1 and PRMT5 genes to their own null mutants (Fig. **6h**). The integration of the HA-tagged PRMT1 and PRMT5 genes to both alleles of their own loci was confirmed by multiplex PCR (Fig. **6h**), and the expression of these proteins was observed by western blotting (Fig. **6i**). Although a significant difference remained between the parental line and the Δ or addback lines, both the percentage of infected THP-1 cells and the number of amastigotes per infected cell were restored upon restoration of PRMT1 and PRMT5 expression (Fig. **6f** and **6g**). In addition, PRMT1, PRMT3 and PRMT5 KO parasites were submitted to a single infection assay with bone marrow-derived macrophages affecting similarly infection (BMDMs, Fig. S3).

**Figure 6.**
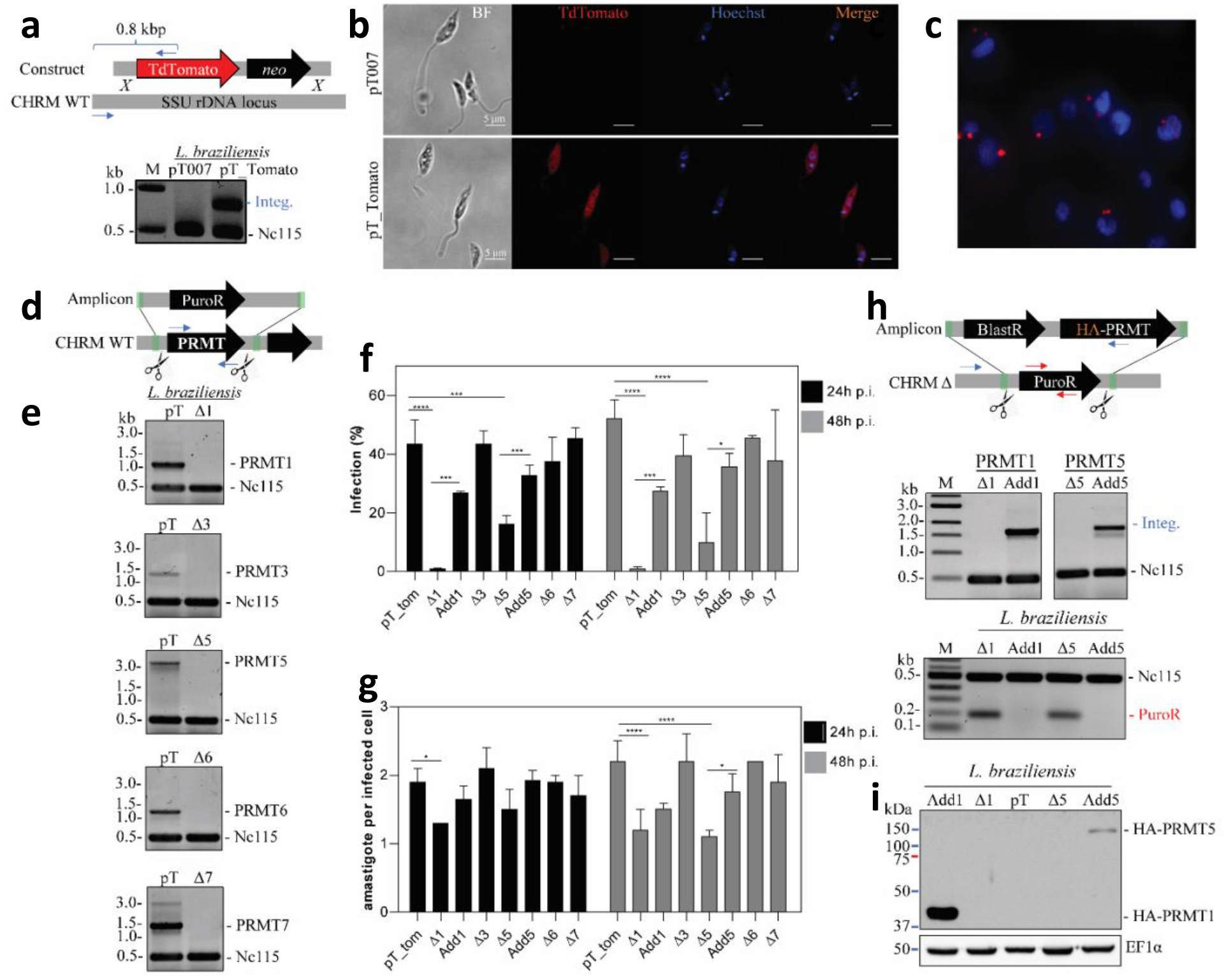
PRMT1- and PRMT5-knockout parasites fail to efficiently infect human macrophages *in vitro*. **a**. Generation of *L. braziliensis* expressing Cas9 (pT) and Tdtomato (tom). Upper panel: schematic representation of the strategy used to integrate the Td_tomato gene into the SSU rDNA locus of *L. braziliensis* pT007. Blue arrows represent primers used to confirm the integration (integ.) of the Td_tomato gene into the SSU rDNA locus; Lower panel: gel electrophoresis of the product of the diagnostic multiplex PCR to confirm Tdtomato integration into the ribosomal rRNA genomic locus (Integ.). **b**. Confocal fluorescence microscopy of the pT007 and pT_tomato parasites. **c**. Representative image of an *in vitro* infection of THP-1 cells with the pT_tomato strain. The images were acquired by ImageXpress Micro XLS System ®, as described in the methods section. **d**. Strategy to knockout PRMTs using pT_tomato parasites; blue arrows represent primers used to detect the presence of PRMT CDSs. **e**. Gel electrophoresis of the products of the diagnostic multiplex PCR of the null mutants. **f** and **g**. *In vitro* infection of THP-1 macrophages evaluated by assessing the percentage of infected cells and the mean number of amastigotes per infected cell. **h**. Generation of addback transfectants. Upper panel: schematic representation of the strategy used to integrate HA-tagged PRMT1 and PRMT5 into the former loci. Lower panels: Gel electrophoresis of the products of the diagnostic multiplex PCR of addback transfectants. Blue arrows indicate primers used to confirm the integration of HA-PRMT1 and 5; red arrows indicate primers used to detect the presence of PuroR. **i**. Expression of PRMT1 and PRMT5 in the addback transfectants (western blotting). Nc115: unrelated gene used as a loading control of the multiplex PCRs (**a, d** and **h**). CHRM WT: wild-type chromosome. Experiments were performed in biological triplicates. \*\**p*<0.002, \*\*\**p*<0.0002 and ****<u>*p*<0.0001</u>, ANOVA Dunnett’s test.

Given the differences observed between the parental and PRMT-KO lines regarding the intracellular survival of amastigotes, we sought to understand whether this was due to impaired resistance of Δ1 and Δ5 mutants to the antimicrobial macrophage machinery or due to impaired appropriate amastigogenesis or amastigote proliferation. We, thus, applied an axenic amastigogenesis strategy in which the differentiating parasites are not subjected to the phagosomal stress conditions (32), and measured the rate of parasite growth by MTT assay (Fig. 7**a**). Interestingly, although equivalent numbers of metacyclic promastigotes were purified from each PRMT-KO line and the parental pT_Tomato line (Fig. 7**b**), the differentiation of metacyclic promastigotes into amastigotes seemed to be impaired in the Δ1, Δ3 and Δ5 parasites, as suggested by an *in vitro* analysis (Fig. **7c**). Concordantly, the number of amastigotes counted at the fourth day of *in vitro* differentiation were significantly reduced in these three mutants (Fig. 7**c**). The re-expression of each of these proteins in their respective null mutants restored their growth and survival to the same rates observed in the parental line (Fig 7**c**). Nevertheless, all the cell lines were able to survive being cultured at 33 °C for two passages in FBS (Fig. 7**a**) and could differentiate back into promastigotes (Supplementary Fig. S4).

**Figure 7.**
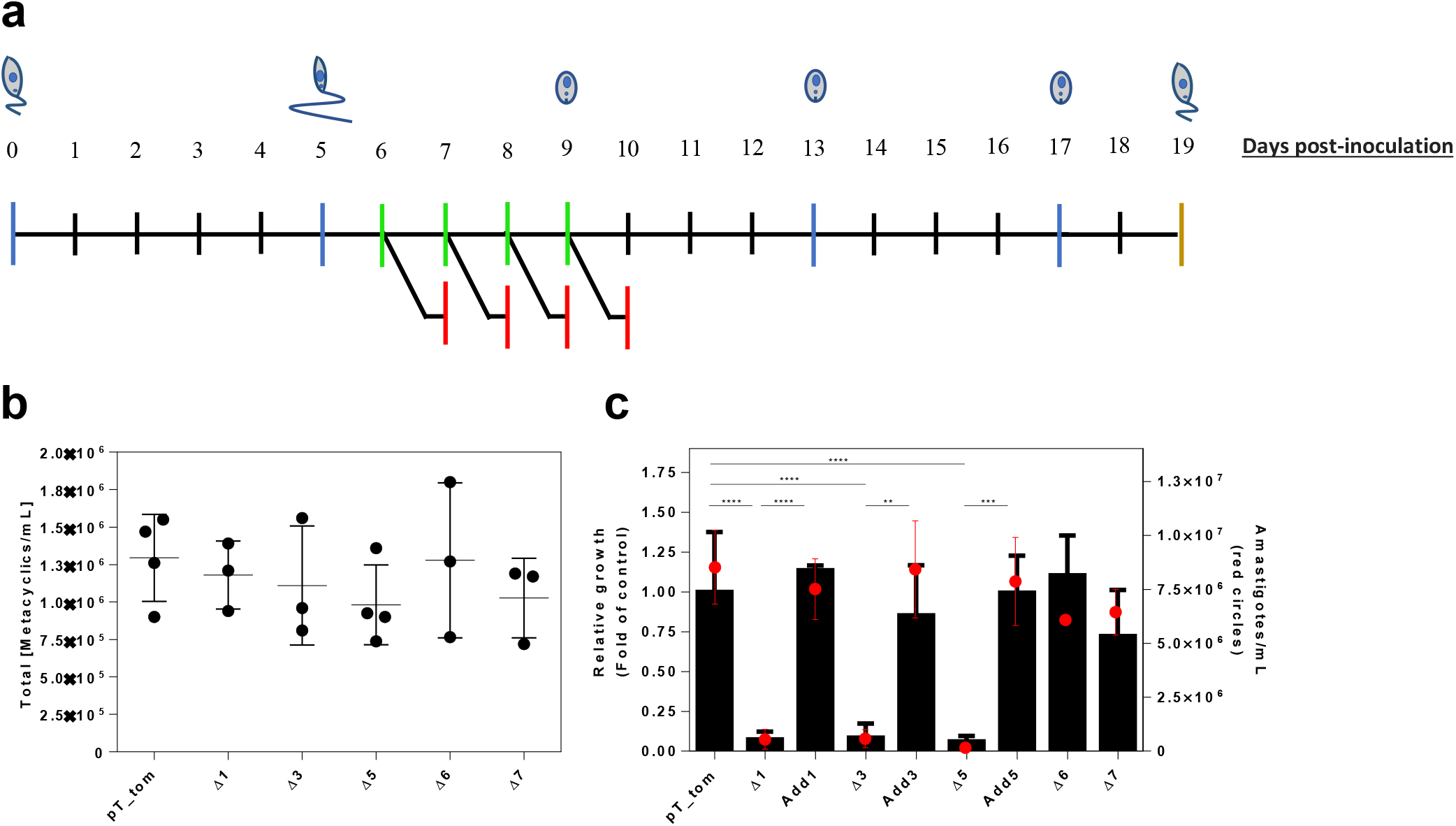
Knockout of LbPRMT1, LbPRMT3 and LbPRMT5 impairs *Leishmania braziliensis* axenic amastigote proliferation *in vitro*. A) Schematic representation of the experimental design to assess the viability and proliferation of PRMT-null mutants of axenic amastigotes *in vitro* compared with pT_Tomato parental cells (WT). On the fifth day post-inoculum, metacyclic promastigotes were isolated by Ficoll gradient, counted and transferred to fetal bovine serum to undergo amastigogenesis. Parasites were collected every 24 h for 4 consecutive days (green lines) and incubated in medium containing MTT for 24 h to determine cell viability (red lines). Amastigotes were subjected to two passages in FBS. At the end of the 12^th^ day of culture, the amastigotes were transferred into complete medium to evaluate their ability to generate viable procyclic forms (yellow line). B) Concentration of metacyclic promastigotes at day 5 post-inoculum. C) The absorbance values after MTT conversion were used in a linear regression (utilizing GraphPad Prism 6) to determine the relative growth of each cell line (*n*≥3). The slope of each line was used as a measure of growth and viability. At the end of the fourth day of differentiation, the number of amastigotes was determined (red circles) using a Neubauer chamber (*n*≥2). **P<0.01, ***P<0.005, ****P<0.001 by one-way ANOVA followed by Tukey’s test.

## 3. Discussion

In this study, we characterized all the canonical protein arginine methyltransferases (PRMTs) in *L. braziliensis*. Among the five annotated *L. braziliensis* PRMTs (*Lb*PRMTs), only *Lb*PRMT3 lacks motifs characteristic of enzymatically active PRMTs. Notably, the global arginine methylation profiles of parasites are developmentally regulated as the parasite progresses from the procyclic promastigote stage to the metacyclic promastigote stage and the amastigote stage. This R-methylation profile also varies among the different *Leishmania* species evaluated. Most importantly, it is evident from this work and previous reports (22, 27, 28) that parasite-derived arginine methylation plays a role in host-parasite interactions. Our data indicate that the expression of major type I and type II PRMTs (i.e., PRMT1 and PRMT5) is required for amastigote survival inside macrophages. The knockout of *Lb*PRMT1 or *Lb*PRMT5 inhibits axenic amastigote growth, which might partially explain the failure of these parasites to infect macrophages *in vitro*. Nevertheless, none of the PRMTs are essential for promastigote growth in axenic culture (Supplementary Fig. S5).

The analysis of the conserved seFquence signatures in the methyltransferase domain revealed the presence of the previously identified key motifs for PRMT activity (6). The PRMT catalytic core is similar across all type I, II and III enzymes, and six signature motifs are relevant for methyltransferase function. Motif I contains three highly conserved glycines and forms the core of the AdoMet binding pocket; postmotif I and motif II are associated with noncovalent bonds and AdoMet binding pocket stabilization; key residues in the double-E motif position arginine residues in substrates; and finally, motif III forms a parallel β-sheet with motif II, and the THW loop is critical for substrate binding and for N-terminal α-helix stabilization. An extra motif, YFxxY, is conserved and located at the N-terminus of Type I PRMTs but absent from type II and III PRMTs (6).

Thus, the alignment of these motifs from *L. braziliensis* PRMTs with the thoroughly investigated motifs from *T. brucei* and *Homo sapiens* PRMTs highlighted the conservation of all six motifs. As expected, the conservation among the trypanosomatid sequences is higher, and it is notable that among the motifs, motif III is less conserved in the 5 sequences that were compared. In contrast to the other PRMTs, both *Lb*PRMT3 and *Tb*PRMT1^PRO^ harbor critical amino acid substitutions in the THW motif (Fig. **1**). However, *Lb*PRMT3 harbors two conserved glutamic acid residues that are essential for the function of the Double E loop motif, while *Tb*PRMT1^PRO^ does not.

Kafková et al. (2017) demonstrated that the enzyme previously named *Tb*PRMT3 is not an active PRMT but acts as a *Tb*PRMT1-like prozyme; therefore, it is currently named *Tb*PRMT1^PRO^. This protein forms a heterodimer with *Tb*PRMT1 (currently known as *Tb*PRMT1^ENZ^), and two heterodimers associate to form the active enzyme unit (21). Although *Lb*PRMT3 possesses a conserved double E loop, the conservation of other sequence features between *Lb*PRMT3 and *Tb*PRMT1^PRO^ provides the first evidence that *Lb*PRMT3 might be, in fact, a prozyme; however, it is still too early to name this enzyme *Lb*PRMT1^PRO^.

In addition, we aligned the five PRMTs from several trypanosomatids and a member of the Bodonidae family, *Bodo saltans*. The alignment results reveal clear subgroups of conservation levels and that the PRMTs from trypanosomes and *Leishmania* cluster together. Interestingly, the results also show that, despite divergences, the PRMT signature motifs are present in the enzymes of the nonparasitic earlier branching *Bodo saltans* species; one exception is the lack of the YFxxY motif in the PRMT6 ortholog, which may impair putative PRMT6 type I activity (Fig. **S1**).

Interestingly, Δ1 and Δ3 mutants displayed a marked reduction in the levels of ADMA, while displaying increased levels of SDMA, and a particularly marked increase of MMA levels (Fig. 4c – 4e). These data indicate that, similar to *T. brucei* PRMT1 (25), *L. braziliensis* PRMT1 is the major type I PRMT expressed in the parasite. As PRMT1 activity was lost in Δ1 and Δ3 cells, the rate of consumption of MMA decreased, causing an accumulation of monomethylated arginine, which in turn increased the availability of monomethylated substrates for type II PRMT activity, leading to an elevation in SDMA levels. This is reinforced by the increased intracellular levels of ADMA, and consequent decrease in MMA and SDMA, upon the re-expression of PRMT1 and PRMT3 (Fig.4g-4i). The individual KO of any other PRMT showed subtle or imperceptible alterations in R-methylation profiles, including the KO of PRMT7, which is proposed to be the only type III PRMT expressed in trypanosomatids. Alternatively, PRMT7 might be responsible for a compensatory increase in activity when PRMT1 or PRMT3 expression is knocked out. Furthermore, we evaluated a group of combined PRMT knockout lines: knockout of two PRMT-encoding genes was performed to reveal the minimum set of classic PRMTs required for *Leishmania* survival. Interestingly, we observed that parasites with both PRMT1 and PRMT3, PRMT5, PRMT6 or PRMT7 knocked out were viable in axenic promastigote culture. Interestingly, Δ1/7 cells displayed a striking reduction in the levels of MMA, showing that, similar to what was observed in *T. brucei* (25), *Lb*PRMT7 is the major catalyzer of MMA formation. The minor changes observed in the MMA profile of the Δ7 cells are likely due to the scavenging of monomethylation activities by other PRMTs. Furthermore, we could not observe any relevant alteration in the levels of arginine methylation upon the knockout of *Lb*PRMT5, even though its ortholog in *T. brucei* (*Tb*PRMT5) is the only type II PRMT characterized in that parasite, and it methylates a relatively broad range of substrates *in vitro* (18). These results may suggest that PRMTs are not at all essential for parasite survival in axenic culture, or these results may indicate redundancy between R-methylation writers and possible noncanonical PRMTs that have not yet been discovered in the *Leishmania* genome.

The expression levels of all the PRMTs vary throughout the *Leishmania* life cycle. PRMT expression is more abundant in promastigotes than in amastigotes, but in particular, the expression of PRMT5 and PRMT7 is lower in amastigotes (8% and 1% procyclic expression, respectively), and the latter is practically undetectable by western blotting (Fig. **3e**). We have also shown that *Lb*PRMTs are predominantly cytoplasmic, except for *Lb*PRMT6, which is more abundant in the nucleus. *Lb*PRMT1, *Lb*PRMT3 and *Lb*PRMT5 show dual nuclear-cytoplasmic localization. Considering that the major classes of arginine-methylated proteins are histones and RNA-binding proteins, it is expected that at least one of the PRMTs should be localized to the nuclear compartment. Since we only analyzed the subcellular localization of PRMTs in procyclic promastigotes, further investigation is necessary to draw conclusions about their localization in other stages of the life cycle.

Using the HA-tagged version of each *L. braziliensis* PRMT, immunoprecipitation assays were conducted to reveal possible targets of these proteins. A large number of proteins were pulled down (135), and we observed that 41% of the interacting proteins (55) were shared among the different PRMTs, leading us to speculate that the parasite has developed a certain redundancy in R-methylation; this redundancy might indicate the relevance of R-methylation for vital processes and at least partially explain why no individual PRMT is required for promastigote viability *in vitro*. Such overlap between PRMTs has been demonstrated in different organisms, including the close relative of *L. braziliensis, T. brucei* (25).

The IP assays also revealed that each PRMT has a variable group of unique interacting proteins (Table S1). In *L. braziliensis*, a large number of proteins were identified in the interactome, 12 of which were PRMT5-specific, 19 of which were PRMT1-specific, 6 of which were PRMT6 specific, and 16 of which were PRMT7 specific. Interestingly, PRMT3 IP pulled down only 2 unique proteins, supporting the hypothesis that PRMT3 does not have R-methylation activity on its own but possibly acts as a cofactor of PRMT1. These unique interacting proteins are of particular interest and should be further explored, especially those of *Lb*PRMT1 and *Lb*PRMT5, which are putative targets for antiparasitic chemotherapy, given their likely role in the parasite-host cell interaction. Importantly, our IP assays revealed an important number of RBPs, which points to the participation of *L. braziliensis* PRMTs as epigenetic regulators of gene expression control pathways in this parasite.

It is intriguing that even with a preferential expression profile in the insect forms, *in vitro* infection is negatively affected when *Lb*PRMT1 or *Lb*PRMT5 are deleted, and our results indicate that *Lb*PRMT1 expression is more abundant in amastigotes than *Lb*PRMT5. It is unknown whether the levels of these proteins correlate with their activities *in vivo*; thus, even when present at very low or undetectable levels, these PRMTs might be able to methylate their targets in amastigotes. Therefore, the R-methylated proteins detected with anti-MMA and anti-SDMA or anti-ADMA antibodies (Fig. **2f, 2g** and **2h**) could have been methylated in one of the previous developmental stages. Importantly, no arginine demethylase function has been reported in this parasite. In fact, we have previously observed that the PRMT7 levels in *L. major* markedly affect not only the rate of cell infection *in vitro* but also the outcome of murine infection *in vivo*, even though the *Lmj*PRMT7 protein levels are largely decreased in the infectious stages of the life cycle (27, 28).

An interesting feature of *T. brucei* PRMT1 is its unique heterotetrameric nature; this complex is composed of two heterodimers containing an active PRMT, *Tb*PRMT1^ENZ^, and a cofactor, *Tb*PRMT1^PRO^. Herein, we did not conduct any functional study to confirm or refute a similar organization of the PRMT1 complex. Nevertheless, we have indications that this is also the case in *L. braziliensis*. We observed that (i) both PRMT1- and PRMT3-knockout *Leishmania* mutants (Δ1 and Δ3) exhibited similarly affected R-methylation profiles, that is, increased MMA and SDMA levels and decreased ADMA levels (Fig. **4c, 4d** and **4e**), (ii) Δ1 parasites had decreased levels of PRMT3 and vice versa (Fig. **5f**); (iii) both proteins exhibited similar localization in procyclic promastigotes (Fig. **3d** and **3f**); (iv) only two PRMT3-specific interacting proteins were pulled down in the co-IP assays (Fig. **5b**), and (v) PRMT3 lacked a conserved TWH motif.

Despite these features, knockout of the PRMT3 gene did not affect the infection profile in the *in vitro* infection assays, unlike knockout of the PRMT1 gene (Fig. **6f** and **6g**). A possible hypothesis would be that *Lb*PRMT1 may play a putative moonlight activity, unrelated to its R-methylation activity, affecting macrophage infection. This hypothesis will be pursued further in future studies. Contrastingly, PRMT3-knockout parasites exhibited impaired axenic amastigote growth at levels similar to PRMT1 knockout parasites, which may be due to the absence of the host-derived factors that play a role in amastigote differentiation and proliferation *in vivo*.

Our data show that there is clear functional overlap among the different *Lb*PRMTs and possible redundancy in their activities. Reverse genetic tools revealed that none of the PRMTs are essential for promastigote axenic growth, but further studies in sand flies would be an interesting future direction. This is supported by the finding that PRMT1- and PRMT5-knockout metacyclic promastigotes are impaired during infection *in vitro* and have reduced abilities to undergo amastigogenesis. In addition, we opened a new venue of investigation by reporting co-immunoprecipitated putative *Lb*PRMT targets, particularly for PRMT1 and PRMT5. Several PRMT inhibitors are currently available for research, and some are already being assessed as anticancer therapies. Based on the phenotypic changes in *Lb*PRMT null mutants described here, inhibition of specific PRMT activity has the therapeutic potential to hinder parasite development in the infected mammalian host and should be tested as antiparasitic drug candidates. A chemical approach that targets *Leishmania* PRMTs may provide feasible ways to block key regulators of the parasite’s gene expression and indirectly affect RNA metabolism.

## 4. Methods

### Alignments

The protein sequences of the PRMTs from trypanosomatids and *Bodo saltans* were obtained from TriTrypDB (https://tritrypdb.org/tritrypdb/app), and the human PRMT sequences were obtained from NCBI (https://www.ncbi.nlm.nih.gov/). The sequences were aligned using Geneious v6.18 software (ClustalW – ID parameters). The key motifs of the conserved PRMT methyltransferase domain are shown. Colors indicate the conservation of residues: the darker the residues are, the more similar they are.

### Parasite cultures

Promastigotes of *Leishmania braziliensis* strain M2903 (MHOM/BR/75/M2903) were cultured in M199 medium (Sigma–Aldrich) supplemented with 10% heat-inactivated fetal bovine serum as previously described (33). To obtain axenic amastigotes, 5×10^6^ promastigotes in the stationary growth phase (fifth day) were inoculated into 10 mL of 100% fetal bovine serum and incubated at 33 °C in 5% CO_2_ as previously described (34). Strains from other *Leishmania* species used were *L. major* LV39 (MRHO/SU/59/P), *L. amazonensis* PH8 (IFLA/BR/67/PH8) and *L. infantum* NCL (MHOM/BR/1972/LD).

### Western blotting

*Leishmania* cells were pelleted (7 min, 3 °C, 2000 xg), washed with 500 µL of ice-cold PBS containing 1X cOmplete protease inhibitors (Roche), resuspended in ∼10 µL extraction buffer (SDS 2%, 50 mM Tris-Cl pH 7,4, 1 mM PMSF, 1x cOmplete) per 1×10^6^ cells and boiled for 10 min. The protein concentrations of the samples were quantified by measuring the absorbance at 280 nm on a Nanodrop One instrument (Thermo Scientific), and the samples were mixed with 0.2 V of 6X sample buffer (350 mM Tris-Cl pH 6.8, 30% glycerol, 10% SDS, 0.12 mg/mL bromophenol blue, 6% 2-mercaptoethanol) and boiled for 3 more minutes. Approximately 40 µg of protein extract was loaded per well in polyacrylamide gels. The proteins were transferred to nitrocellulose membranes (GE Healthcare Life Sciences – REF 10600003), which were incubated with the appropriate antibodies according to the manufacturer’s instructions. Anti-MME (Cell Signaling Technologies [CST] – 8015S), anti-SME2 (CST – 13222), anti-AME2 (CST – 13522), anti-HA (Sigma – H3663), and anti-EF1α (Merck – 05-235) were used for western blotting according to the manufacturer’s instructions. Secondary anti-mouse and anti-rabbit peroxidase-conjugated antibodies (GE Healthcare – NA931V and NA934V, respectively) were used according to the manufacturer’s instructions. The membranes were visualized using the chemiluminescent substrate of an ECL kit (GE Healthcare – RPN2232). Images were captured on ImageQuant LAS 4000 (GE Healthcare).

### Single guide and donor DNA construction

The protocol for the construction of sgRNA templates was slightly modified from Beneke et al., 2017. For the amplification of sgRNA templates, 0.2 mM dNTPs, 2 µM each of primer G00 (sgRNA scaffold) and a gene-specific forward primer (primers 29–41 – Table S2), and 1 unit of Phusion DNA Polymerase (NEB) were mixed in 1× HF buffer in a total volume of 50 µl. The PCR conditions were 30 s at 98 °C followed by 35 cycles of 10 s at 98 °C, 30 s at 55 °C, and 15 s at 72 °C. A second PCR was carried out using 1 µl of the abovementioned reaction as the template for amplification with G00_loredited_F and G00_loredited_R (same conditions; 50 µl total volume) to obtain higher yields. For donor DNA amplification, specific primers (primers 42–51 and 14–18 in Table S2) were used to amplify the resistance genes from pTBlast or pTPuro (35) and 3xHA_Nterm_Blast (sequence 65 – Table S2). The PCR conditions were 30 s at 98 °C followed by 35 cycles of 10 s at 98 °C, 40 s at 60 °C, and 2 min at 72 °C (2 reactions of 50 µl for each donor). All the reactions described above were combined and precipitated using ethanol and sodium acetate. The DNA pellet was resuspended in 60 µl of 1x Tb-BSF buffer and used to transfect parasites in the log phase of growth. To generate the Addback transfectants (Fig. **4f-i**), the PRMT1 and PRMT3 genes were amplified with primers 54–57 (Table S2) and cloned between the *Xho*I and *Bam*HI sites of pSSU_neo_Tdtomato (36). The resultant plasmids were digested with *Pac*I and *Pme*I (New England Biolabs) and used to transfect *L. braziliensis* Δ1 and Δ3.

### Transfections

Promastigotes in the log phase of growth were centrifuged at 2000 g at 25 °C for 10 minutes. The pellets were resuspended in 1/10 V *Tb*-BSF buffer (37). The parasites were centrifuged under the same conditions and resuspended at a concentration of 2×10^8^ parasites per ml in the same buffer. One hundred microliters of this suspension was mixed with the proper amount of DNA, and transfection was performed with an Amaxa Nucleofactor instrument (LONZA) using the X-001 program. The culture was maintained in M199 for 16 hours prior to further selection with the appropriate antibiotics (6 µg/ml G418; 16 µg/ml hygromycin B; 10 µg/ml blasticidin; 10 µg/ml puromycin) on solid M199 media.

### Immunofluorescence

A total of 1.5×10^6^ promastigotes in the log phase of growth were fixed with 3% paraformaldehyde for 20 min at room temperature. The fixed cells were adhered to poly-D-lysine (Millipore)-coated glass slides and permeabilized with 0.2% Triton X-100 for 5 min. The cells were blocked with 5% skim powder milk in TBS-T for ½ h. An anti-HA (Sigma H3663) primary antibody was diluted in 2.5% milk with TBS-T at 1:500 and incubated with the samples for 2 h. Secondary antibodies conjugated to Alexa Fluor 488 (Invitrogen A11001) at a 1:500 dilution in TBS were added and incubated for ½ h. Hoechst (Invitrogen H3570) at a concentration of 2 µg/ml in TBS was used to stain the nuclei and kinetoplasts for 15 minutes. Images were acquired with an Axio Observer combined with an LSM 780 confocal device at 630× magnification (Carl Zeiss). Images were processed with Fiji ImageJ free software.

### Immunoprecipitation

A total of 5×10^8^ parasites in the log phase of growth were centrifuged (2000 g, 10 min, 4 °C), and the pellets were washed with 20 ml of ice-cold PBS and centrifuged. The resulting pellets were resuspended in 1 ml of lysis buffer (Tris-Cl pH 8.0 20 mM, NaCl 150 mM, Triton X-100 0.5%, EDTA 0.5 mM, cOmplete mini [Roche] 3X), frozen in N_2_ and thawed on ice. A 21G needle was used to resuspend the suspension 20 times on ice to lyse the parasites, and 4 U of Turbo DNAse was added to each sample. The samples were centrifuged (10000 g, 10 min, 4 °C), and the pellets were discarded. The supernatants were incubated with 30 µl of protein A/G-coated magnetic beads (Genscript L00277) to remove the nonspecifically bound proteins for 1 hour at 4 °C with agitation. Ten micrograms of specific antibodies (anti-HA [Sigma H3663] or anti-MME [CST - 8015S]) were diluted in 500 µl of lysis buffer and incubated with 100 µl of the beads for 1 hour at room temperature with agitation. The unbound antibodies were removed using a magnet, and the beads were incubated with precleared supernatants for 2 hours at 4 °C with agitation. The beads were washed 6 times with 1 ml of 0.05% TBS Tween using a magnet, and the bound proteins were eluted by boiling the beads with 60 µl of sample buffer 2X (117 mM Tris-Cl pH 6.8, 10% glycerol, 3.3% SDS, 0.04 mg/mL bromophenol blue, 2% 2-mercaptoethanol) for 5 minutes. A fraction of the eluted proteins were used for western blotting, and the remaining proteins were sent to the University of Laval for mass spectrometry identification.

### Proteomic analysis

The data obtained by mass spectrometry were analyzed using Scaffold 4 software (http://www.proteomesoftware.com/products/scaffold/). In the case of the anti-HA IP, the proteins identified in the WT sample (negative control) with a ≥50% probability of being correctly identified were removed from the analysis. After this removal, the criterion for identifying proteins that interacted with each PRMT was ≥95% protein identification probability. Interactivenn (38) was used to generate a Venn diagram. Tools of TriTrypDB (https://tritrypdb.org/tritrypdb/app) were used to search for proteins among the PRMT interacting proteins that harbor RGG motifs or gene ontology terms related to RNA binding. For the MME IP analysis, proteins whose normalized total spectra were at least 2 times higher in Δ1 were listed.

### THP-1 culture and differentiation

THP-1 cells were cultured in complete RPMI-1640 medium (Sigma–Aldrich) supplemented with 10% heat-inactivated fetal bovine serum (Sigma–Aldrich) and 0.5% penicillin-streptomycin (Thermo Fisher Scientific) and incubated at 37°C in a 5% CO_2_ atmosphere. The cells were not allowed to exceed a maximum density of 10^6^ · cells mL^-1^. For cell differentiation, THP-1 monocytes were incubated with 30 ng·mL^-1^ 4α-phorbol 12-myristate 13-acetate – PMA (Sigma Aldrich) for 24 hours (39). After the differentiation period, the medium with nondifferentiated cells was removed, and the cells were incubated in fresh RPMI-1640 medium for 48 hours for complete maturation and improvement of phagocytic properties (40); then, the cells were infected with *Leishmania*.

### *In vitro* infection assay

The infection assays were performed at 33 °C in a 5% CO_2_ atmosphere using differentiated THP-1 macrophage-like host cells seeded in 96-well black flat-bottom plates at 2 10^4^ cells·mL^-1^. After the macrophage differentiation and maturation period, the THP-1 differentiated macrophage-like cells were infected with metacyclic promastigotes (10 parasites:1 host cell) purified during the stationary growth phase by Ficoll density gradient (41). After 4 hours of incubation, noninternalized parasites were removed by washing the cells with RPMI-1640 medium, and the plates were incubated at 33 °C in a 5% CO_2_ atmosphere in fresh medium with 0.25% Hoechst (Merck-Millipore) to stain the DNA. At 24 and 48 hours postinfection, *Leishmania* intracellular growth was monitored by analyzing fluorescent images acquired by the ImageXpress Micro XLS System® (Molecular Devices, LLC, EUA); 9 fields of view of each well plate were examined under 40x magnification. DAPI filters (λ_ex_: 377 and λ_em_: 447) and Texas Red filters (λ_ex_: 562 and λ_em_: 624) were used to image the cell nuclei and *Leishmania* amastigotes, respectively. The cell and parasite counts were obtained using the algorithm Transfluor from MetaXpress Software (Version 5.3.0.5, Molecular Devices, LLC) by analysis of the acquired images. To count the nuclei, the algorithm was set to detect structures between 3–9 µm and intensities 1600 gray levels above background, while structures between 2.5–5 µm and intensities 1700 gray levels above background were used to count *Leishmania* (vesicles). The infection rate (%) was calculated based on the ratio of infected cells to the total THP-1 cells counted, and statistical analysis (Dunnett method) was performed to identify significant differences in infection and/or growth between the control and knockout parasites.

For BMDM infection, bone marrow was harvested from BALB/c mouse femurs and tibias by cutting both ends of the bones and flushing out the marrow with PBS. Precursor cells were cultured in complete RPMI supplemented with 30% L929 cell-conditioned medium. After 7 days in culture, the mature BMDMs were harvested by washing the cells with cold PBS, incubating on ice for 15 min and pipetting up and down several times. One day prior to infection, 1 × 10^5^ cells were plated on 13-mm diameter coverslips and incubated at 37 °C in 5% CO_2_ for 16 h in complete RPMI. The cells were infected with metacyclic promastigotes at an MOI of 1:1 for 4 h and washed with incomplete RPMI to remove free parasites. The cells were incubated in fresh complete RPMI for 24 h and 72 h. Coverslips were collected at each time point, stained using the Diff‐Quick method (LaborClin, Brazil) and mounted onto glass slides. The slides were analyzed under light microscopy, and 100 macrophages in each of the replicates were counted on each coverslip.

### Evaluation of amastigote growth in culture

*Leishmania braziliensis* M2903 pT_Tomato parental cells or PRMT-null mutants were seeded at 5× 10^5^ cells/mL in 10 ml of complete M199 medium and incubated at 26 °C for five days until they reached the stationary phase of growth. On the fifth day post inoculum, metacyclic promastigotes were isolated by Ficoll gradient (41) and counted using a Neubauer chamber. Metacyclic promastigotes were transferred to 10 ml of fetal bovine serum (FBS) at 2× 10^5^ cells/ml and incubated in vented flasks in a humid atmosphere at 33 °C in 5% CO_2_ to undergo amastigogenesis. To verify the cell viability of the different PRMT-KO cell lines during the process of amastigogenesis, two milliliters of each culture was harvested every 24 h for 4 consecutive days, centrifuged at 1,300 xg for 10 min, washed once with PBS and resuspended in 1 ml of M199 containing 1 mg/ml MTT ((3-(4,5-dimethylthiazol 2-yl)-2,5-diphenyltetrazolium bromide). Next, 200 µl of the cell suspensions were added to each well of 96-well plates (4 wells per cell line) and incubated at 26 °C for 24 h. The rate of MTT conversion (i.e., the increase in absorbance at 570 nm) was used in a linear regression in GraphPad Prism 6, and the slope of the line was used as a measure of the relative amastigote growth in culture. At the end of the fourth day of culture in FBS, 500 µl of the fully differentiated amastigotes were inoculated in 4.5 ml of FBS (first passage) and incubated at 33 °C for four days. Then, the amastigotes were subjected to a second passage in FBS for four more days. At the end of the 12th day, 500 µl of the same culture was inoculated into 4.5 ml of M199 to evaluate for their ability to generate viable procyclic promastigotes. After 48 h of culture in M199, the number of procyclic cells was quantified by counting in a Neubauer chamber.

## Acknowledgements

We thank Tania P. A. Defina for the technical support. We thank Eva Gluenz (University of Glasgow) for providing us with the plasmids for performing CRISPR/Cas9, and we thank Pegine Walrad and Michael Plevin (University of York) for the very productive discussions.

## Ethics statement

All mouse experimental procedures were performed in accordance with the Ethical Principles in Animal Research approved by the Local Ethical Animal Committee (CEUA) of FMRP-USP (protocol 107/2020).

The funders had no role in study design, data collection and analysis, decision to publish, or preparation of the manuscript.

