## Supplementary Figures for "Functional study of *Leishmania braziliensis* protein arginine methyltransferases (PRMTs) reveals that PRMT1 and PRMT5 are required for macrophage infection"

**Supplementary Lorenzon et al., 2021**

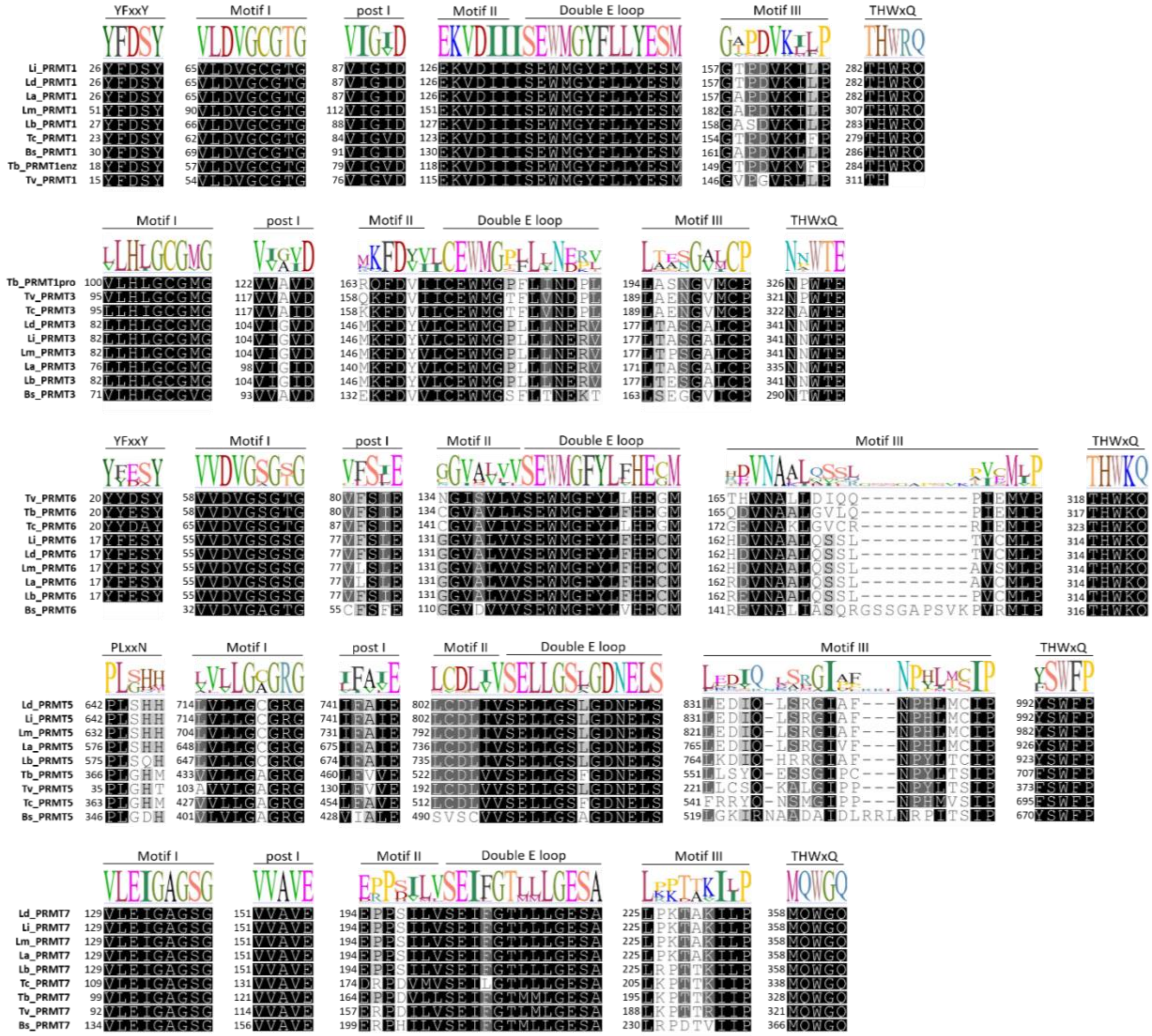


**Figure S1.** Kinetoplastid PRMT orthologs. Alignment of PRMTs from *L. braziliensis*, *L. amazonensis*, *L. major*, *L. donovani*, *L. infantum*, *T. cruzi, T. brucei*, *T. vivax* and *Bodo saltans.* The sequences of proteins were aligned using Geneious v.6.18 software. The regions corresponding to key motifs of PRMT activity are represented as a consensus logo.

Addback PCR

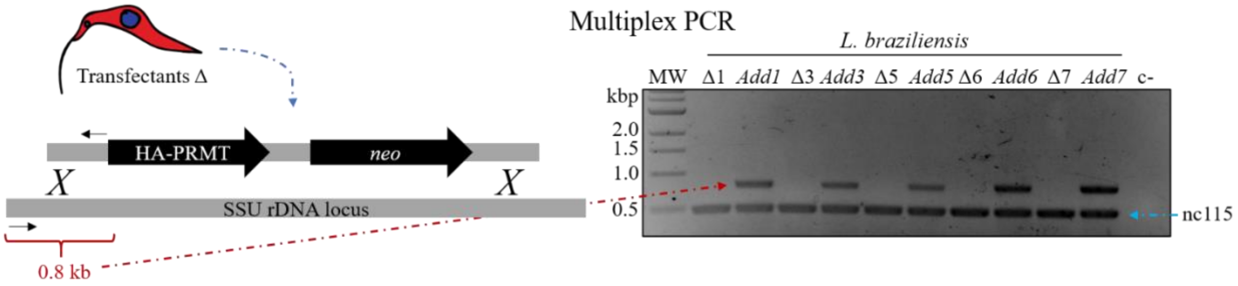


**Figure S2**. Generation of Addback parasites. Multiplex PCR using primers represented by black arrows in the cartoon (primers 63 and 64 – table S2) and primers that amplify the unrelated nc115 gene (primers 52 and 53 – table S2).

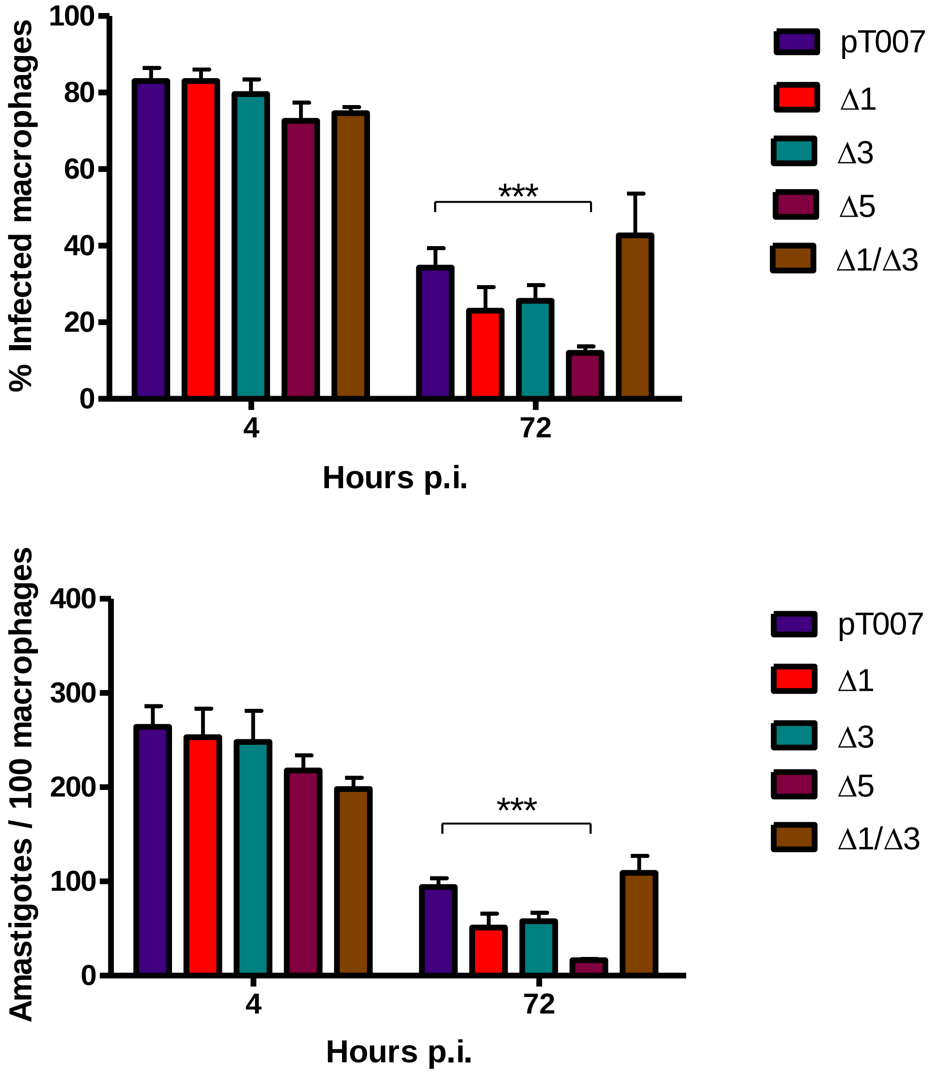


**Figure S3**. In vitro infection of Bone Marrow Derived Macrophages with PRMT null mutants.

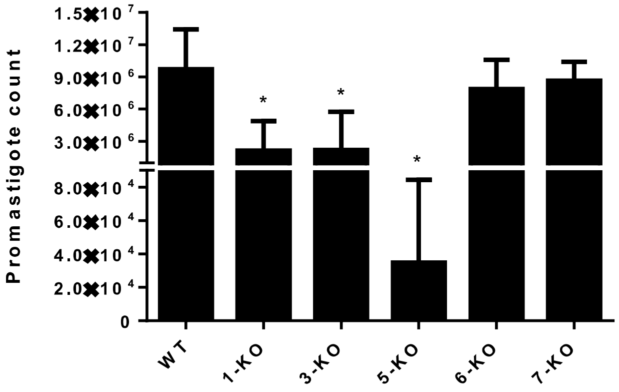


**Fig. S4.** Knockout of *Leishmania braziliensis* PRMTs does not impair amastigote-to-promastigote differentiation *in vitro*. After two passages in FBS, amastigotes were transferred into complete M199 and incubated at 26°C. After 48h, the number of promastigotes originated was determined for each cell line in three independent experiments (except for PRMT5-KO, n = 2) using a Neubauer chamber. *P<0.05, ****P<0.001 by one-way ANOVA followed by Dunnett's test.

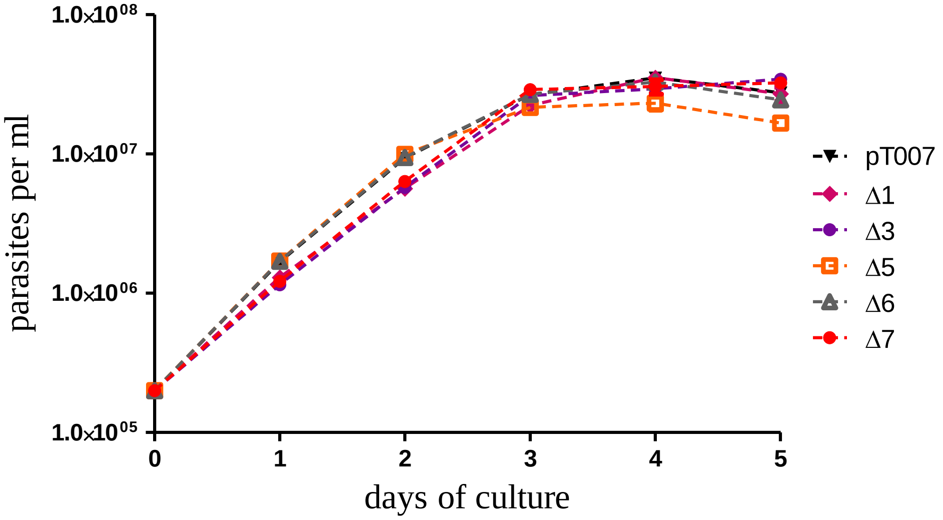


**Figure S5.** Growth curve of PRMT null mutants. Promastigotes were inoculated at $2x10^5$ cells per mL in triplicate and counted every 24 hours in Neubauer chamber for 5 days. No statistically significant difference was observed between the null mutants and parental line pT007.
